# MolEvolvR: A web-app for characterizing proteins using molecular evolution and phylogeny

**DOI:** 10.1101/2022.02.18.461833

**Authors:** Faisal S Alquaddoomi, Joseph T Burke, Lo Sosinski, David A Mayer, Evan P Brenner, Samuel Z Chen, Jacob D Krol, Ethan P Wolfe, Vince P Rubinetti, Shaddai Amolitos, Kellen M Reason, John B Johnston, Janani Ravi

## Abstract

Studying proteins through the lens of evolution can reveal features such as conserved domains, lineage-specific variants, and co-occurring domain architectures in phylogenetic context across all superkingdoms. *MolEvolvR* enables researchers to conduct such evolution-focused studies to generate testable hypotheses about protein function and evolution. *MolEvolvR* is a novel web-app allowing researchers to visualize the molecular evolution of their proteins of interest in a phylogenetic context across the tree of life. It accepts multiple input formats – protein/domain sequences, homologous proteins, or domain scans – and, using a general-purpose computational workflow, returns detailed homolog data and dynamic graphical summaries (e.g., phylogenetic trees, multiple sequence alignments, domain architectures, domain proximity networks, phyletic spreads, co-occurrence patterns across lineages). *MolEvolvR* performs domain-centric searches to capture remote homologs that are missed by full-length searches, integrates domain architecture evolution with phyletic distribution analyses, and provides evolutionary context visualizations that reveal lineage-specific adaptations versus those that are broadly conserved. Thus, *MolEvolvR* is a powerful, easy-to-use web interface for computational protein characterization. The web-app can be accessed here: https://jravilab.org/molevolvr.

---

The rate at which protein families are being discovered far outpaces that at which molecular or biochemical functions are assigned to proteins^1^. This gap impedes scientific understanding of critical cellular processes such as molecular pathogenesis and stress response. Identifying and characterizing the complete molecular systems involved in such processes requires functional knowledge of the proteins involved. Numerous studies^2–18^ have demonstrated the power of phylogenetic and molecular evolutionary analysis in determining the molecular functions of such proteins. A variety of individual tools^19–45^ exist for protein characterization, including sequence similarity searches or ortholog detection, delineating co-occurring domains (domain architectures), and building multiple sequence alignments and phylogenetic trees. However, these tools operate in isolation, requiring users to manually integrate and interpret results within an evolutionary context. Despite providing complementary insight, these tools are disjointed and often require both technical skill and background knowledge for meaningful use. Biologists need a means to functionally characterize their unknown/understudied protein(s) of interest by exhaustively identifying relevant homologs with shared motifs, domains, and domain architectures. Moreover, homology searches using full-length protein sequences often fail to identify remote homologs that share only individual functional domains.

Here, we present *MolEvolvR*, a unified, user-friendly, and accessible web-based framework that addresses these limitations by utilizing data from all taxonomic domains to analyze biological sequences of interest in depth (**Fig. 1**; accessible at jravilab.org/molevolvr). Given one or more proteins of interest from the user, *MolEvolvR* performs both protein- and domain-centric searches to capture remote homologs, reconstructs domain architecture evolution across the tree of life, and provides evolutionary context visualizations that distinguish broadly conserved features from lineage-specific adaptations. These findings, spanning multiple taxonomic scales, are effectively summarized and visualized to generate functional hypotheses. The computational evolutionary framework underlying *MolEvolvR* is written using custom R^46–61^ and shell scripts. The web application is built with an R/Shiny^48,62,63^ framework with additional user interface customizations in HTML, JavaScript, and CSS, and has been tested on Chrome, Firefox, and Safari browsers on Mac, Windows, and Linux operating systems.

**Figure 1.**
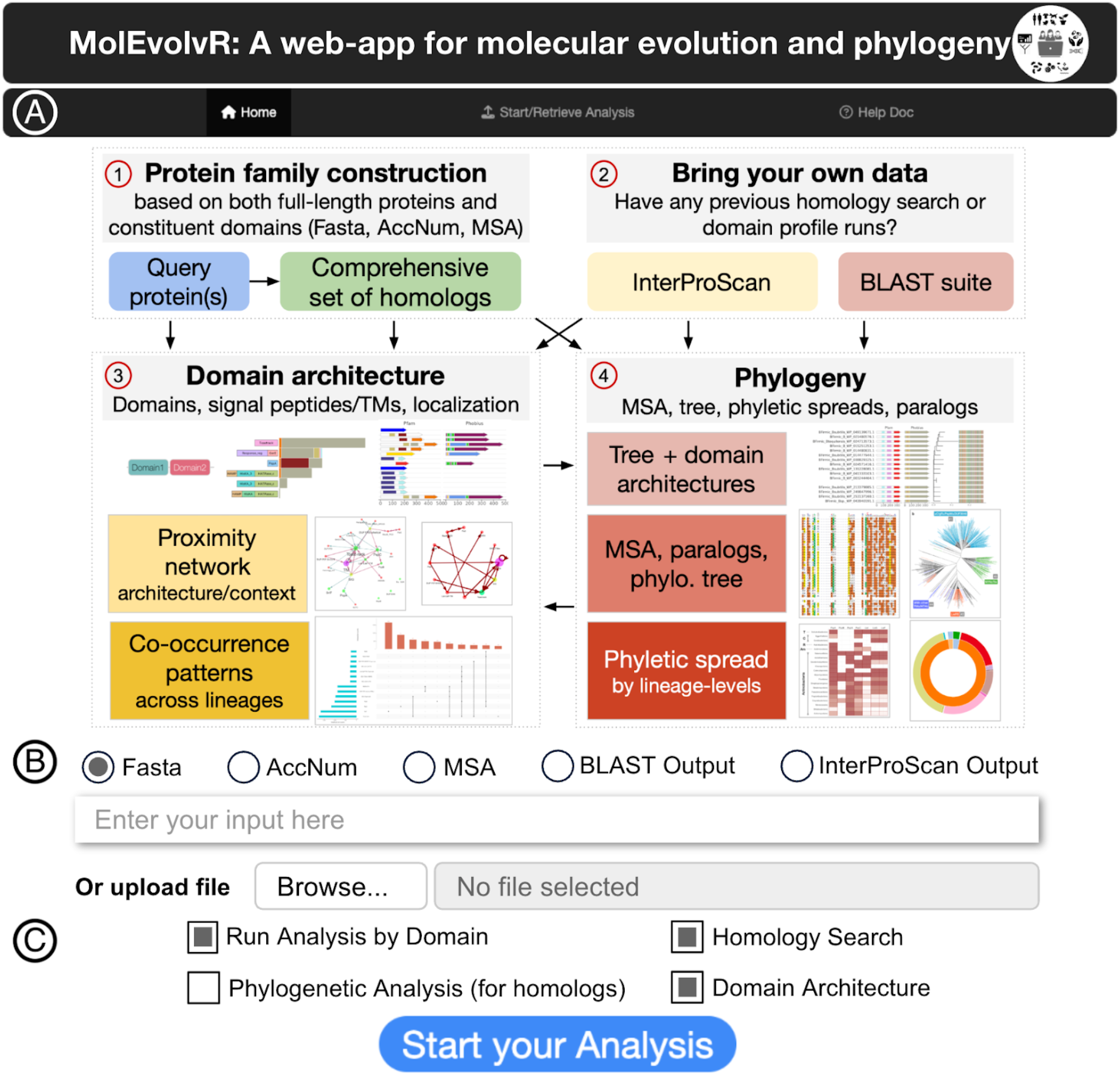
Overview of *MolEvolvR*. **A**. *MolEvolvR* allows users to start with protein(s) of interest and perform the complete analysis (1+3+4), only protein characterization (1+3), or only homology searches (1+4); or start with external outputs from BLAST or InterProScan for further analysis, summarization, and visualization (2+3+4). *MolEvolvR* is interactive, queryable, and customizable. **B**. Multiple input options in *MolEvolvR*: FASTA, NCBI/UniProt accession numbers, pre-computed multiple sequence alignment, MSA (in FASTA formats), analysis outputs from an external web or command-line BLAST or InterProScan runs. **C**. The different analysis options available in *MolEvolvR* depend on the chosen input formats in **1A, 1B**. See *Supplementary case study* and web-app help docs (https://jravilab.org/molevolvr/?r=&p=help) for further details on versatile input/output options and data summarizations and visualizations.

Consider a researcher starting with a protein of unknown function, either identified through a genetic screen for a phenotype or an uncharacterized protein in a genomic segment of interest [**Fig. S1**]. In Step 1, *MolEvolvR* resolves this query protein into its constituent domains and uses the protein and each of its domains for iterative homology searches^20,21^ across the tree of life^64–67^ [**Fig. S2**]. This divide-and-conquer strategy is domain-sensitive by design and captures remote homologs missed by homology searches using full-length protein sequences alone. In Step 2, *MolEvolvR* characterizes the protein and each of its homologs by reconstructing their domain architectures and delineating putative molecular function [**Fig. S3**], combining: 1) sequence alignment and clustering algorithms for domain detection^19,24,25,68^; 2) profile matching against protein domain and orthology databases^32,33,69–72^; and 3) prediction algorithms for signal peptides^33,73^, transmembrane regions^33,74–76^, cellular localization^33,75^, and secondary/tertiary structures^33,77–80^. In Step 3, *MolEvolvR* maps each of these homologs and their domain architectures to their lineages to further tease out broadly conserved vs. phylum-specific signatures, placing these findings in the context of evolution [**Fig. S4**]. This analysis illustrates the breadth of *MolEvolvR*’s analytic capabilities: the results entail a detailed molecular characterization of the initial query protein(s) and their homologs, integrated across the superkingdoms of life, revealing both universally conserved features and any lineage-specific functional variants.

Beyond single protein inputs, *MolEvolvR* is versatile, accommodating a variety of inputs for investigating wide-ranging questions and producing diverse output summaries and visualizations in easy-to-access formats. Queries can include protein/domain sequences of single- or multi-protein operons (FASTA, NCBI/UniProt protein accession numbers), homologous proteins (pre-computed MSA or web/command-line BLAST output), or motif/domain scans (InterProScan output) [**Fig. 1B**]. *MolEvolvR* tailors analyses to answer a variety of questions, e.g., determining protein or functional domain features restricted to certain pathogenic groups using the organism filter to discover virulence factors [**Fig. 1C**; **Fig. S1**]. Finally, *MolEvolvR* generates output types spanning a complete set of homologs or phylogenetic trees, the domain architectures of a query protein, or the most common partner domains [previews in **Fig. 1A**]. Along with tables and visualizations, *MolEvolvR* also provides a dashboard with summary statistics and graphical summaries integrating these results in the context of evolution [**Fig. S1**]: i) structure-based multiple sequence alignments and phylogenetic trees; ii) domain proximity networks from all co-occurring domains (across homolog domain architectures) consolidating within and across query proteins; iii) phyletic spreads of homologs and their domain architectures; and iv) co-occurrence patterns and relative occurrences of domain architectures across lineages [**Fig. 1A**]. The domain-centric searches trace the evolution of the proteins/domains of interest, even across remote homologs. The web-app contains detailed documentation about all these options, including case studies (see *Supplementary Material*) and frequently asked questions (FAQs). In addition to easy access to a local, in-browser history of user-submitted jobs, *MolEvolvR* can, optionally, send users a detailed description and status update of their submitted jobs via email. Finally, all figures, tables, and reports can be downloaded by the user.

*MolEvolvR*’s integrative approach has enabled the discovery of functional properties across diverse protein systems. Analysis of the phage shock protein (PSP) stress response system identified ∼20,000 homologs and ∼200 novel domain architectures, revealing evolutionary variants adapted to different membrane stress challenges across bacteria, archaea, and eukaryotes^2^. Domain-centric searches captured homologs sharing core functional domains (e.g., PspA) despite <30% overall sequence identity, while phyletic spread analyses distinguished universally conserved components from lineage-specific adaptations. A specific instance and adaptation of *MolEvolvR*, applied to delineate the PSP stress response system in great detail across the tree of life, is available at https://jravilab.org/psp^2^ [see *Supplementary Case Study*]. *MolEvolvR* is a generalized web server of this web-app. To demonstrate its broad applicability, we have successfully applied the approach underlying *MolEvolvR* to discover and functionally characterize several systems, including proteins/operons in zoonotic pathogens, e.g., two nutrient acquisition systems in *Staphylococcus aureus*^4,5^, a novel phage defense system in *Vibrio cholerae* (informing functional validation experiments)^6^ surface layer proteins in *Bacillus anthracis*^7^, helicase operators in bacteria^8^, CARD^81^ antimicrobial resistance genes and uncharacterized proteins associated with AMR by our machine learning models^82^, and internalins in *Listeria* spp^9^. We have included a few pre-loaded examples in *MolEvolvR* for users to explore, in addition to help pages and FAQ. Researchers can submit requests for new analysis or visualization features through *MolEvolvR*.

Thus, *MolEvolvR* (jravilab.org/molevolvr) is a flexible, user-friendly, and powerful interactive web tool that enables researchers of any level of bioinformatics experience to bring molecular evolution and phylogeny to bear on their proteins of interest and provide evolutionary context and domain organization that inform functional hypotheses and guide experimental validation.

## Acknowledgments

We would like to thank members of the JRaviLab for testing the web-app at various stages and providing the authors with several iterations of constructive feedback. We are also profoundly grateful to Krishnan Raghunathan, Arjun Krishnan, Emily Meyer, Jerome McKay, Premal Shah, Dave Bunten, and members of the Krishnan lab (Kayla Johnson and Nathaniel Hawkins) for their early feedback on *MolEvolvR* and detailed comments on the manuscript. We have benefited from several diverse use cases and challenges brought to us by our collaborators that helped fine-tune the functionality of the web-app (inline citations), and the larger *MolEvolvR* user community that has engaged with us to keep the app current, relevant, and feature-rich.

## Author Contributions

JR conceived and designed the study; JTB, LS, SZC, JDK, EPB, and JR acquired the data and performed all the analyses; FSA, JTB, LS, EPB, DAM, JDK, JR, FSA, and DAM made the figures and tables; FSA, JTB, LS, DAM, SZC, JDK, JR wrote the backend code; FSA, JTB, DAM, SZC, JDK, VPR, JR built the R/Shiny web-app; FSA, DAM, JBJ, KMR, and SA set up the server that hosts the web-app; FSA, EPB, DAM, EPW, JTB, VPR, JDK, LS, JR tested the web-app rigorously; EPB, EPW, FSA, DAM, JTB, JDK, LS, JR worked on the help documentation; LS and JR wrote the first draft of the manuscript; EPB, JTB, JDK, LS, EPW, and JR revised the manuscript.

## Data Availability and Reuse

All the data, analyses, and visualizations are available in our interactive and queryable web application: http://jravilab.org/molevolvr. The web-app is licensed under the MIT license.

