## Supplementary material for "MolEvolvR: A web-app for characterizing proteins using molecular evolution and phylogeny"

#### Supplementary Text: *A Case Study*

Here, we present a case study with the *lia* operon (from *Bacillus subtilis*) involved in lantibiotic stress response, a well-studied phage shock protein (PSP) stress response system[2]. The PSP systems are known for combining conserved components (e.g., PspA) and variant themes (e.g., centering on PspBC or Lia components). It is initially unclear what underlying building blocks (domains/domain architectures) might culminate in the stress response function and their history in the broader context of bacterial evolution (phylogeny, phyletic spreads). Therefore, this section focuses on a detailed characterization of the six Lia proteins, LiaIHGFSR; the detailed results (analyzed data and graphical summaries) are available here for interactive exploration: jravilab.org/molevolvr/?r=liasix (see "liadom" *MolEvolvR* code for domain-wise analyses of the *lia* operon). We used *MolEvolvR* to conduct a comprehensive analysis across the three superkingdoms of life and delineate all occurrences of Lia proteins in terms of their underlying domain architectures and phyletic spreads. The full report can be downloaded from the 'Results Summary' tab.

***Homologs*.** We first generated the homologs for each of these six proteins across all completed representative or reference genomes (from bacteria, archaea, and eukaryota) [Data tab, web-app]. The homolog data from these genomes are available in an interactive and queryable format in the *MolEvolvR* web-app with linked NCBI accession numbers, species, lineages, percentage similarities, and protein- and similarity-search-related parameters [**Fig. S2**]. Best hits by species or lineage can be easily subsetted, and the entire data table is available for download in a standard CSV format.

***Domain architectures*.** We next determine the domain architectures (based on sequence-structure motifs/domains, disorder predictions, transmembrane regions, signal peptides, and cellular localizations) of the six Lia proteins and each of their homologs. In the 'Domain Architecture' tab, we summarize our results for individual and across all query proteins [**Fig. S3A**, left]. In addition to the diverse domain architectures, we can also determine their phyletic spreads using a simple pop-up feature to determine widespread vs. lineage-specific domain architectures [**Fig. S3A**, right]. Using *MolEvolvR*, we can also visualize the diverse domain architectures and cellular localizations (e.g., Pfam, Gene3D, Phobius, MobiDB) of representative homologs of each Lia protein to discover both similar and dissimilar homologs (e.g., novel partner domains, variants with altered localization [**Fig. S3B**]). We summarize our domain findings by profile database/prediction algorithm in the 'Network' section of the 'Domain Architecture' tab [**Fig. S3D**]. The proximity network consolidates the findings across proteins (or by protein) by connecting co-occurring domains within proteins by their frequency of occurrence across lineages (nodes = domains; edges = co-occurrence; node size and width of arrows = frequencies of occurrence). This big-picture view, along with the co-occurrence plots [**Fig. S3E**] helps discover new components and their relative frequencies across thousands of homologs. This approach also leads to serendipitous connections across homologs of the different Lia proteins (e.g., two-component LiaRS system variants that carry both the response regulator and histidine kinase domains), as well as predominantly lone domains (e.g., PspA_IM30) [**Fig. S3E**].

***Phylogeny.*** Finally, we used *MolEvolvR* to study the evolution of the Lia proteins [Phylogeny tab, *MolEvolvR*]. We recorded the presence of homologs, by lineage, through interactive sunburst plots [**Fig. S4A**; Phylogeny tab, web-app] and heatmap for all query proteins [**Fig. S4B**; Data tab, web-app]. We generated multiple sequence alignments and phylogenetic trees based on key representatives from diverse species, lineages, and domain architectures [**Fig. S4C–D**, Phylogeny tab, web-app].

Through these comprehensive analyses, we first characterized the proteins encoded by the *lia* operon in terms of their domain architectures, with LiaH, a PspA stress response effector protein, two transmembrane proteins LiaI and LiaG, two globular domains of unknown function with Toastrack-like domains in LiaF and LiaG, and a two-component system LiaRS with response regulator/receiver domain and histidine kinase. We discovered several homologs in lineages outside Firmicutes for each of the Lia proteins and domains, including the widespread two-component system. In contrast, other domains (PspA, DUF2154, DUF4097) were predominantly present in only Firmicutes genomes. We also identified new connections within the more extensive Lia proximity network based on homologs of the LiaRS proteins, with proteins that carried domains from both response regulator and histidine kinase domains (e.g., in Proteobacteria, Planctomycetes), and rare DUF4097-containing homologs within Firmicutes that carry an additional N-terminal DUF1700 or a second DUF4097 domain.

We performed a more comprehensive analysis of the phage shock protein (PSP) system and its partner domains across the tree of life using a *MolEvolvR*-like approach, which led to multiple novel discoveries in a recent article[2]. We report ~20K homologs of key PSP proteins, ~200 novel domain architectures (present in ≥2 species), and ~500 genomic contexts (≥5 species). Due to the variety of domain architectures, cellular localizations, and phyletic spreads of each of these protein families and operons, the PSP system is a well-suited use case to exemplify how we can use molecular evolution and phylogeny to characterize proteins. This approach also contributed to the discovery and functional characterization of the *avcD* phage defense system that we found to be broadly conserved in several divergent bacteria, archaea, and eukaryotes; this finding was further validated by the complementation of bacterial *avcD* function by yeast *avcD* homolog[6]. The PSP work, including this *lia* operon use case and other recent diverse biological applications[2–11], indicates that the *MolEvolvR* approach is invaluable in characterizing new protein families of interest in the context of evolution.

### Supplementary Figures

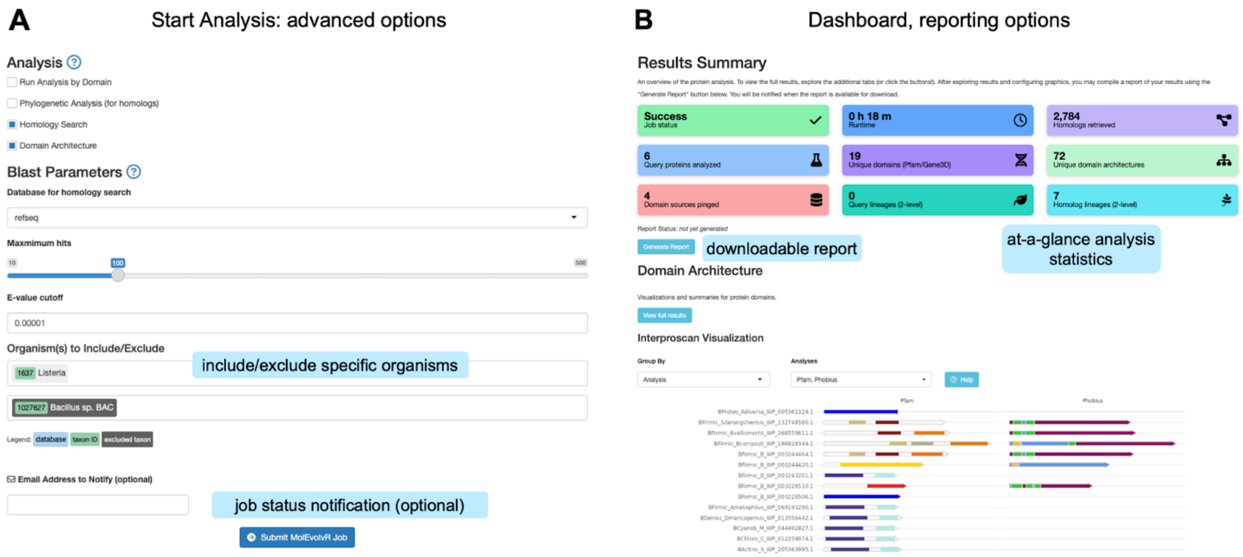


Figure S1. Analysis options, results dashboard, and reports

**“Start Analysis” and “Analysis Results” pages** include parameters for the analysis and results of the analysis, respectively. Panel **A** shows the advanced options available when starting an analysis, including the nature of analysis, selection of reference database, homolog thresholds, job status, and the ability to include and/or exclude specific organisms in the resulting analysis. Panel **B** displays the Results Summary page, which features large tiles that provide an at-a-glance overview of the analysis processing and results. The “Generate Report” button creates a downloadable HTML version of the analysis that can be used offline, shared, or archived. The report includes interactive graphics and tables similar to those displayed on the *MolEvolvR* site. *Light blue boxes with text are not part of the screenshot; they have been added to highlight key features for clarity.*

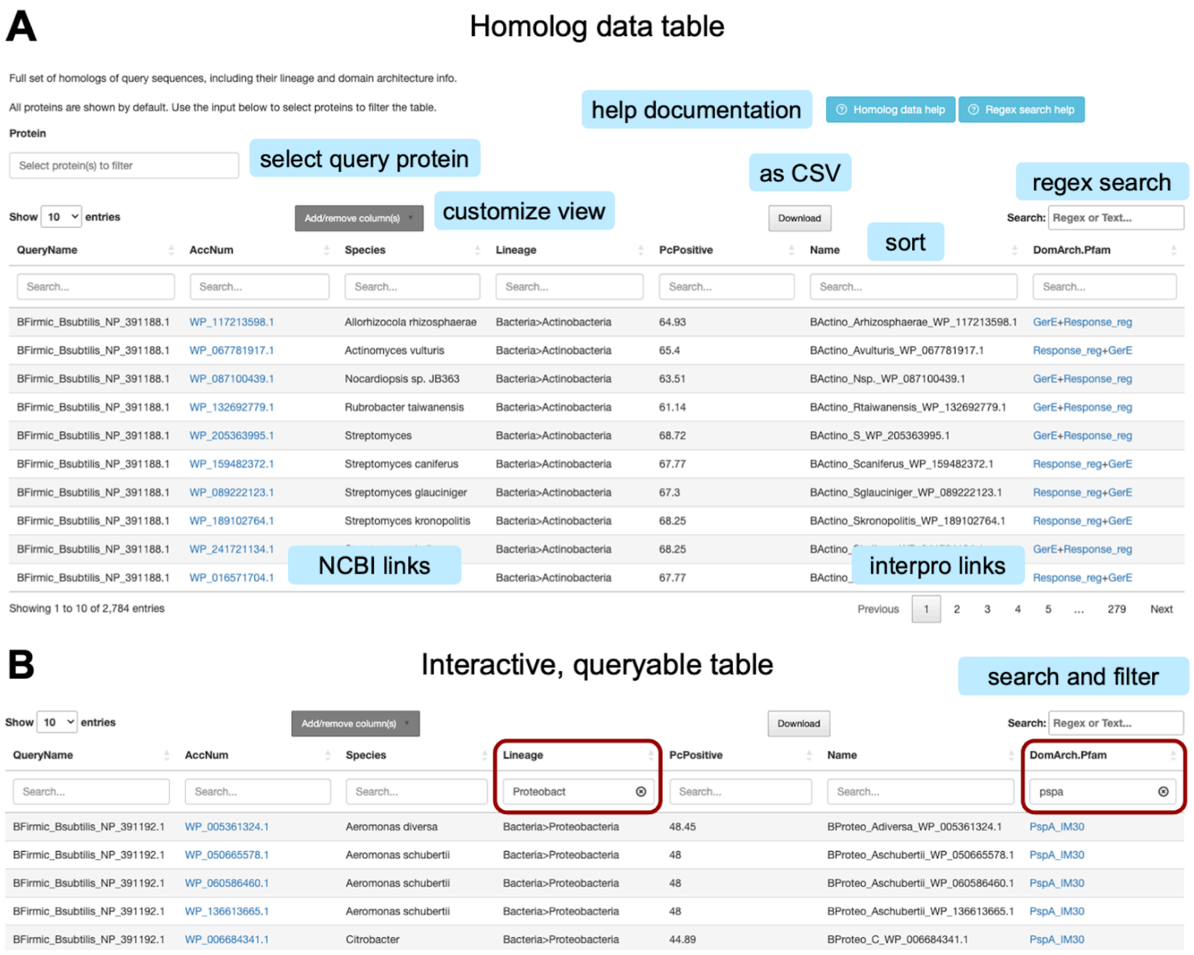


Figure S2. Explore the homologs, their lineages, and domain architectures with MolEvolvR.

**Homolog data table** with the best hits from all superkingdoms of life (queried across all RefSeq genomes). For each homolog, we tabulate details across all query proteins, including genome, species, lineage, sequence homology (BLAST parameters), and domain architecture information in a sortable (enabled for all columns), queryable, and interactive manner. The data updates dynamically, such as when selecting which query protein(s) to view homologs and companion data for (top left). The 'Add/remove column(s)' button allows users to access the complete list of columns and modify them as needed. Results can be subsetted using the search box, which can optionally persist across tabs for fine-tuning each analysis. Additionally, the search supports regular expression (regex) queries. 'Download' allows the user to download either the filtered subset of the table or the entire table for further local, programmatic, or spreadsheet-based analysis (as a CSV file). The accession number for each homolog is hyperlinked to its corresponding NCBI protein page. Domains are hyperlinked to EBI's InterProScan details pages. Panel **A** displays a snapshot of the top hits across all query proteins, using the default columns. Panel **B** shows an example of the **interactive, queryable table** with homologs being filtered based on specific lineages (e.g., Proteobacteria) AND Pfam domain architecture (e.g., PspA). *Light blue boxes with text are not part of the screenshot; they have been added to highlight key features.*

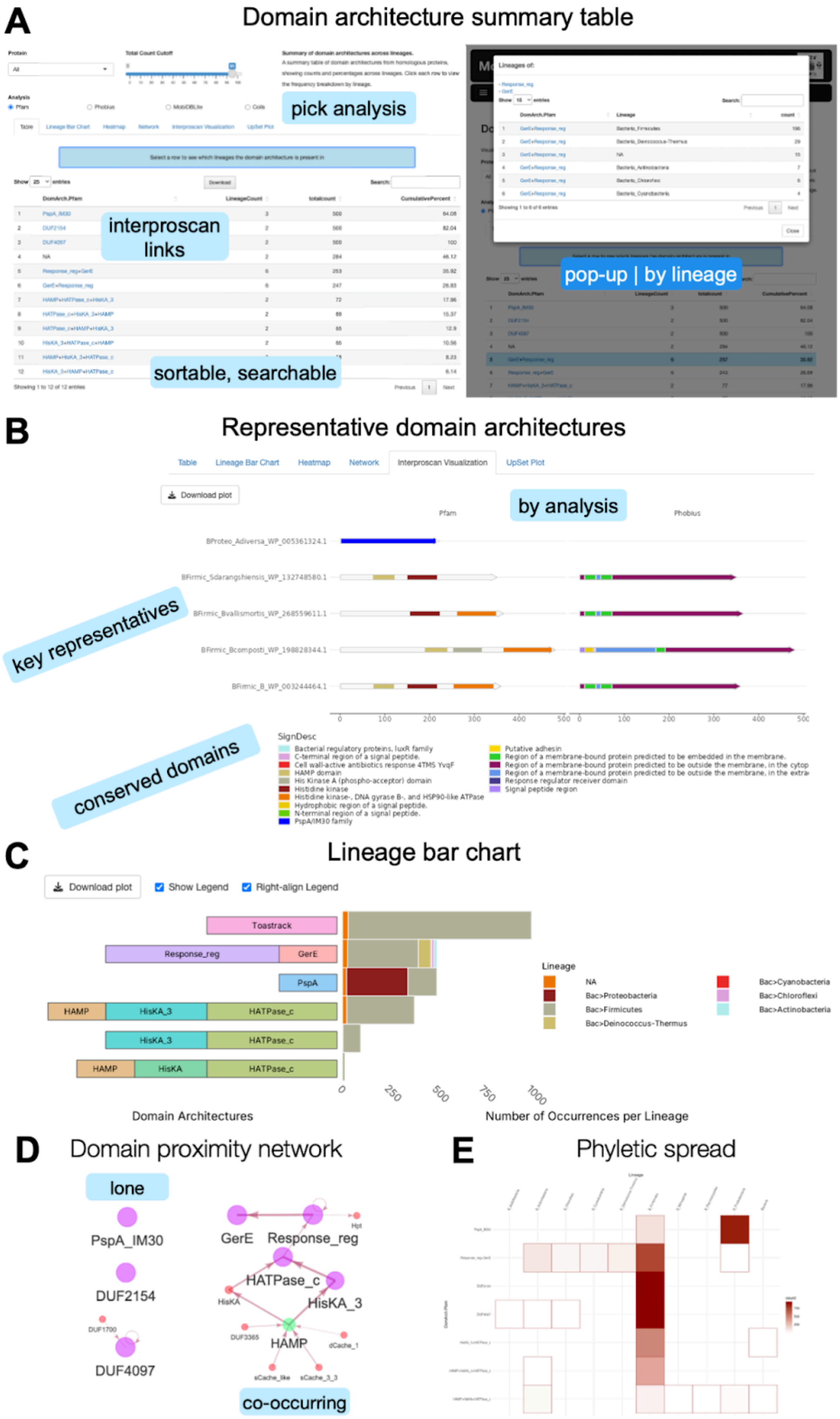
A
Domain architecture summary table
pick analysis
interproscan links
sortable, searchable
pop-up | by lineage
B
Representative domain architectures
Table
Lineage Bar Chart
Heatmap
Network
Interproscan Visualization
UpSet Plot
Download plot
by analysis
Pfam
Phobius
key representatives
conserved domains
SignDesc
C
Lineage bar chart
Download plot
Show Legend
Right-align Legend
Toastrack
Response_reg
GerE
PspA
HAMP
HisKA_3
HATPase_c
HisKA
Lineage
NA
Bac>Proteobacteria
Bac>Firmicutes
Bac>Deinococcus-Thermus
Bac>Cyanobacteria
Bac>Chloroflexi
Bac>Actinobacteria
0
250
500
750
1000
Domain Architectures
Number of Occurrences per Lineage
D
Domain proximity network
lone
PspA_IM30
DUF2154
DUF4097
GerE
Response_reg
HATPase_c
HisKA_3
HAMP
co-occurring
E
Phyletic spread

Figure S3. Domain architectures of query proteins and their homologs using *MolEvolvR*.

**A. Domain architecture summary table** with phyletic spreads. **Left:** The table shows the top (most predominant) domain architecture (or 'domarchs', in the figure) by query protein (or across all queries, as in this case) with the frequency of occurrence and lineages in which they occur. The slider enables users to select the top hits, which represent 98% of all homologs in this case. **Right:** The second snapshot shows the pop-up that appears when each domain architecture row is clicked, elaborating on the 'LineageCount' by displaying the frequencies of occurrence by individual lineage for the selected domain architecture. **B. Representative domain architectures** with one per lineage-domain architecture combination to show the diversity. The figure displays cartoon representations of key representative domain architectures (Pfam column) and cellular localizations (Phobius column) for the query protein homologs (rows). The Pfam and Phobius annotations for each domain prediction (by colour) are shown in the legend. **C. Lineage bar chart.** This graphic shows the top domain architectures and proportions of the primary lineages that carry them. **D. Domain proximity network.** The network captures co-occurring domains within the top 98% of the homologs of all the 'query' Psp members and their key partner domains (after sorting by decreasing frequency of occurrence). The size and colour of the nodes (domains) and the width of edges (co-occurrence of domains within a protein) are proportional to the frequency of their occurrence across homologs. The complete network, as well as networks specific to each query, are available on the web-app. **E. Phyletic spread** of the predominant domain architectures across query proteins. The heatmap shows the presence/absence of homologs of all query proteins across key lineages (columns) for each predominant domain architecture (rows). The colour gradient indicates the highest number of homologs in a particular lineage. The heatmap gives the whole picture. *Rows*: Top domain architectures across all homologs. *Columns*: The major archaeal, bacterial, eukaryotic, and viral lineages with representative sequenced genomes. *Light blue boxes with text are not part of the screenshot; they have been added to highlight key features.*

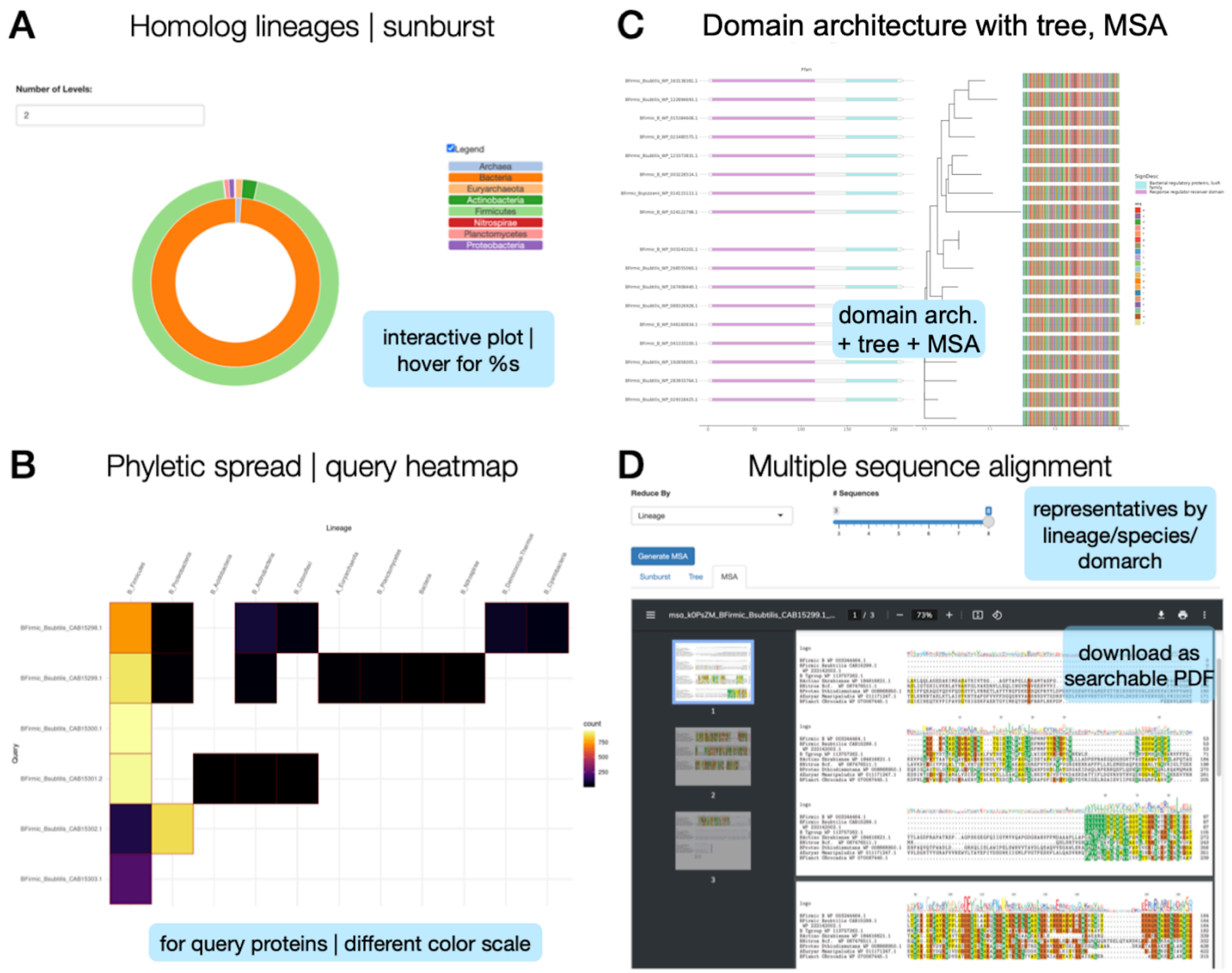


Figure S4. Phyletic spreads and phylogeny of query proteins using *MolEvolvR*.
**A. Lineages of homologs.** The sunburst plot shows the phyletic spread of all the homologs. The legend shows the lineages (inner ring, kingdoms; outer ring, phyla) that carry homologs. Note: The sunburst plots only display lineages of >0.1% fraction of total proteins. **B. Phyletic spread** of the homologs by query protein. The heatmap shows the presence/absence of homologs across key lineages (columns) for each query (rows). The colour gradient indicates the highest number of homologs in a particular lineage. The heatmap provides a comprehensive view across domain architectures and query proteins. *Rows*: Query proteins are queried against all sequenced and completed genomes across the three major superkingdoms (also referred to as Domains of life). *Columns*: The major archaeal, bacterial, eukaryotic, and viral lineages with representative sequenced genomes. **C.** The **multiple sequence alignment** is overlaid on the **phylogenetic tree**. In the tree generated for LiaS (BFirmic_Bsubtilis_CAB15299.1), each leaf (protein) is named with its kingdom (e.g., ‘B’ for bacteria), phylum (first six letters, e.g., ‘actino’ for Actinobacteria), genus, species (represented as ‘Gspecies,’ e.g., ‘Bsubtilis’ for *Bacillus subtilis*), and NCBI protein accession number (e.g., CAB15298.1), resulting in a uniquely identifiable name for the protein, ‘KPhylum_Gspecies_AccNum,’ e.g., BFirmic_Bsubtilis_CAB15299.1. *Key*: the colours in the

multiple sequence alignment depiction correspond to different amino acids. **D.** Snapshot of the **multiple sequence alignment** of representative homologs of LiaS (BFirmic_Bsubtilis_CAB15299.1) by lineage. The generated MSA, with representative homologs from each lineage, species, or domain architectures, is available to users as a downloadable, searchable PDF. *Light blue boxes with text are not part of the screenshot; they have been added to highlight key features.*
